# AMPK-activator ATX-304 reduces oxidative stress and improves MASLD via metabolic switching

**DOI:** 10.1101/2024.02.13.578901

**Authors:** Emanuel Holm, Isabeau Vermeulen, Saba Parween, Ana López-Pérez, Berta Cillero-Pastor, Michiel Vandenbosch, Silvia Remeseiro, Andreas Hörnblad

**Affiliations:** Department of Medical and Translational Biology, Umeå University, Johan Bures väg 12, 90187 Umeå Sweden; Maastricht MultiModal Molecular Imaging Institute (M4i), Maastricht University, Maastricht, Limburg, the Netherlands; The MERLN Institute for Technology-Inspired Regenerative Medicine, Department of cell Biology-Inspired Tissue Engineering, Maastricht University, Maastricht, Limburg, the Netherlands; Wallenberg Centre for Molecular Medicine (WCMM), Umeå University, 90187 Umeå, Sweden

## Abstract

Metabolic dysfunction-Associated Steatotic Liver Disease (MASLD) is the most common chronic liver disease worldwide for which there are no approved treatments. Adenosine monophosphate-activated protein kinase (AMPK) is an interesting therapeutical target since it acts as a central regulator of cellular metabolism. Despite efforts to target the AMPK, no direct activators has yet been approved for treatment of this disease. This study investigates the effect of AMPK activator ATX-304 in a preclinical mouse model of progressive fatty liver disease. The data demonstrate that ATX-304 diminishes body fat mass, lowers blood cholesterol levels, mitigates liver steatosis, and ameliorates the development of liver fibrosis. The beneficial effects of ATX-304 treatment are accompanied by a shift in the liver metabolic program, including increased lipid oxidation, reduced lipid synthesis, as well as remodeling of cholesterol and lipid transport. We also observed variations in lipid distribution among liver lobes in response to ATX-304, and a shift in the zonal distribution of lipid droplets upon treatment. Taken together, our data suggest that ATX-304 holds promise as a potential treatment for Metabolically Associated Fatty Liver Disease (MAFLD), including in human patients.

## Introduction

Metabolic dysfunction associated steatotic liver disease (MASLD, former NAFLD/MAFLD) is the most common chronic liver disease worldwide, affecting approximately one third of the global population (1,2). Metabolic dysfunction steatohepatitis (MASH), the severe form of MASLD, is characterized by hepatic steatosis, inflammation, liver damage and resulting fibrosis. It is one of the leading causes of cirrhosis and end-stage liver disease, including liver cancer. The prevalence of MASH is expected to increase in the coming decade (3), paralleling the global obesity and Type 2 Diabetes (T2D) epidemic, thus becoming an increasingly important risk factor for liver cancer (4).

Despite the progress that has been made in understanding the etiology and pathophysiology of MASLD, there are no drug treatments licensed for the disease. Current improvements rely primarily on lifestyle changes, including diet, exercise, and weight loss. Still, various drugs are under development with the aim to resolve MASH and achieve reduced fibrosis (5,6). AMP-activated protein kinase (AMPK) is an important protein that is involved in regulating whole-body energy homeostasis and that has received much attention as a potential target in MASLD, but also in metabolic diseases in general, including T2D. AMPK is a cellular sensor of energy status that is activated upon low energy supply such as after caloric restriction (7) or exercise (8). It has been shown that activation is stimulated by hormones involved in insulin sensitivity and fatty acid metabolism (9,10), and that some of the positive metabolic effects of the commonly used T2D drug metformin are mediated by indirect activation of AMPK.

Upon activation AMPK signaling adjusts cellular metabolism by inhibiting anabolic processes and promoting catabolic pathways (11). Genetic studies in mice have addressed the function of AMPK in the liver, and in the context of MASLD and MASH (12–17). Evidence show that AMPK inhibits hepatic de novo lipogenesis and cholesterol synthesis (18), and it has also been suggested to stimulate fatty acid oxidation (17). This is to a large extent mediated by inhibition of Acetyl-CoA Carboxylase (ACC) and HMG-CoA Reductase (HMGCR), via phosphorylation (19–23). Reduced activity of AMPK is part of the pathology of MASLD and various AMPK activators have been tested in animal models, corroborating the idea that AMPK targeting holds promise as a potential target for treatment in MASLD and MASH (13,18,24,25). Nevertheless, to date no direct AMPK-activators have yet reached the market, (27).

Recently, the dual AMPK and mitochondrial activator ATX-304 (formerly ATX-304) was shown to improve glucose homeostasis and cardiovascular function in T2D patients on metformin as well as in diet-induced obese mice (28,29). It also reduces insulin resistance and improves cardiac function and ameliorates diabetes in mouse models with impaired beta-cell function (30,31). Given these data and the long-standing interest in the AMPK-pathway for treatment of metabolic disease and specifically MASLD, we investigated the effect of ATX-304-treatment on mice fed a choline-deficient high-fat diet (CD-HFD) that develop progressive liver disease including hepatic steatosis and MASH with resultant fibrosis (32). We performed transcriptomics, proteomics, spatial and bulk lipidomics analysis as well as histological and biochemical characterization of livers from these mice. Our study demonstrates that ATX-304 reduces body fat mass, blood cholesterol and liver steatosis, as well as ameliorates the development of liver fibrosis in CD-HFD mice. The effects of ATX-304 were accompanied by a shift in the metabolic program of the liver including increased fatty acid oxidation, reduced fatty acid synthesis, as well as changes in cholesterol and lipid transport. ATX-304 treatment also reduced the abundance of harmful oxidized lipids in treated livers. In addition, we detected lobular heterogeneities in lipid distribution in response to ATX-304 and a consistent switch in the zonal distribution of lipid droplets upon treatment. Taken together, our data suggest that ATX-304 is a promising candidate for the treatment of MASLD also in human patients.

## Results

### ATX-304 treatment reduce whole-body fat mass and blood cholesterol of male CD-HFD mice

To investigate the potential positive effects of ATX-304 treatment on MASLD and MASH, 5-week-old male C57Bl/6J mice were fed choline-deficient high-fat diet (45% kcal fat) for 3w or 21w and then switched to a CD-HFD formulated with ATX-304 (1 mg/g, CD-HFD+ATX-304). The short-term cohort was treated for 7w before sacrifice while the long-term cohorts were treated for either 10w or 24w (Figure 1A). Control mice were fed a normal chow diet. Ingestion of CD-HFD over a longer period (21w) induced liver steatosis, inflammation, and fibrosis, while the shorter period (3w) represents earlier stages of disease. For all cohorts, food intake and body weight were measured regularly (Figure 1B, Supplementary figure 1A-C). Noticeably, within two weeks from the switch to ATX-304-diet for the long-term experiments, the body weights of treated mice were reduced to levels similar to control mice on a regular diet (Figure 1B). Moreover, EchoMRI measurements of body composition at start and end of the treatment demonstrated that weight reduction corresponds to a loss in fat mass, while lean weight was not reduced (Figure 1C-E). In line with previously published data (28,29), ATX-304 also greatly improved glucose homeostasis and reduced insulin levels in treated mice (Supplementary figure 2).

**Figure 1.**
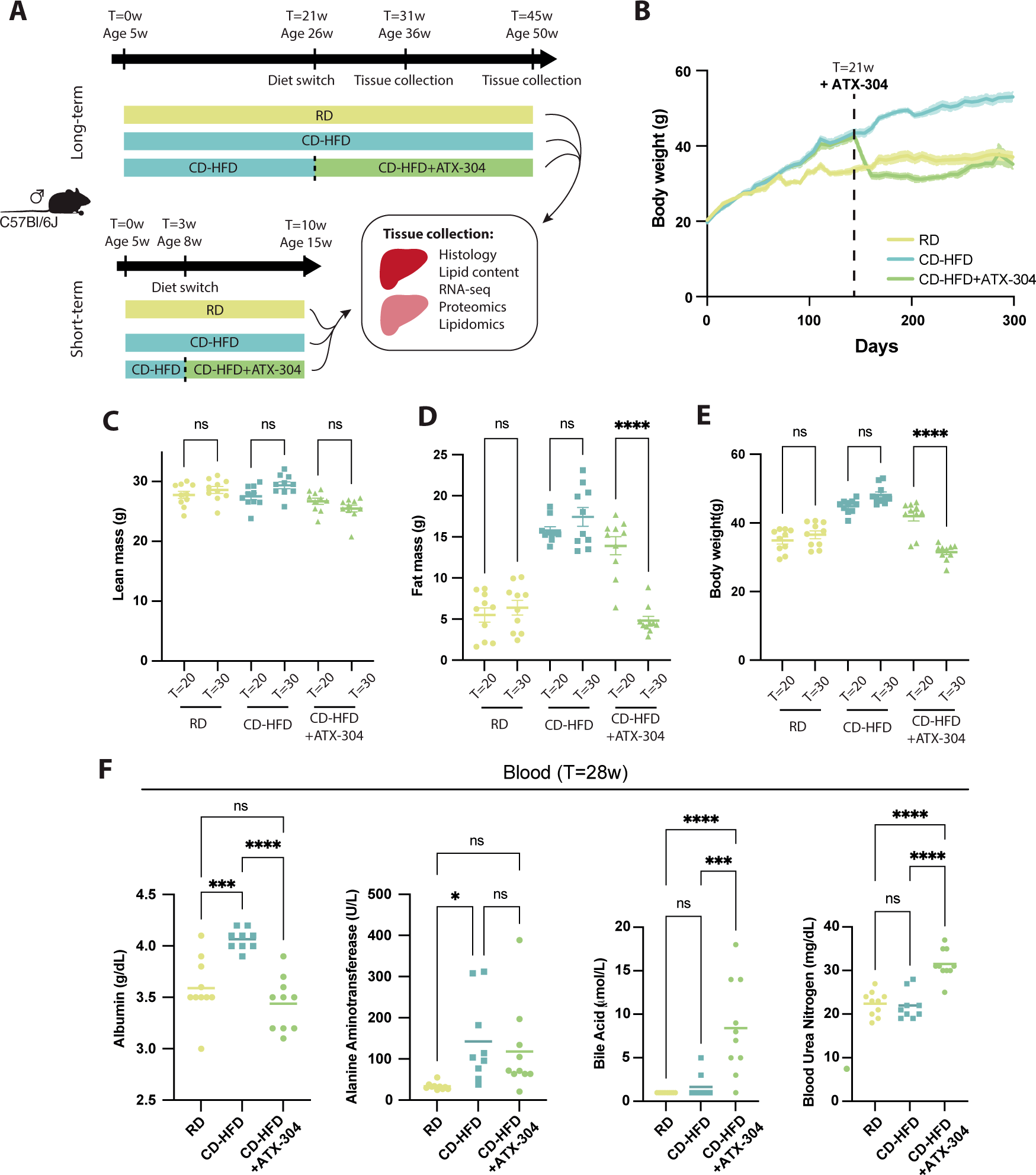
ATX-304 reduce body fat mass and blood cholesterol in male CD-HFD mice. A) Overview of mouse cohorts and experimental workflow. **B**) Weight curve for long-term RD, CD-HFD and CD-HFD+ATX-304 mice. Dashed line indicate start of ATX-304 treatment (T=21w). Colored shade depicts standard error of the mean (SEM). Lean (**C**), fat (**D**) and total body mass (**E**) before (T=20w) and after (T=30w) ATX-304 treatment. ****p<0.0001 (One-way ANOVA with Tukey’s multiple comparisons test) (n=10 for all groups). **F**) Vetscan blood profiling of albumin (left), alanine aminotransferase, bile acid and blood urea nitrogen for RD, CD-HFD and CD-HFD+ATX-304 mice. *p<0.05, ***p<0.001, ****p<0.0001 (One-way ANOVA with Tukey’s multiple comparisons test). Individual data points, mean ± SEM are indicated in all graphs (n=10 for all groups).

### ATX-304 lowers blood cholesterol and alters cholesterol metabolism in the liver

We further assessed liver function using the Vetscan 2 “mammalian liver profile” system 8w after start of treatment (T=28) and found that blood albumin levels were normalized in ATX-304 treated CD-HFD mice (Figure 1F). CD-HFD had significantly higher levels than the RD control mice (Figure 1F). No significant difference in the levels of alanine aminotransferase was detected between ATX-304- treated mice and untreated CD-HFD mice. However, alanine aminotransferase levels of CD-HFD+ATX- 304 mice were not significantly increased compared to that of control RD mice (Figure 1F). In contrast, blood levels of both bile acids and blood urea nitrogen were elevated in ATX-304-treated animals as compared to both RD controls and non-treated CD-HFD mice (Figure 1F). Moreover, blood cholesterol was almost doubled in CD-HFD mice (212 mg/dL) compared to RD control mice (114 mg/dL), while blood cholesterol was restored down to almost normal levels (143 mg/dL) in ATX-304-treated CD-HFD mice (Figure 2A).

**Figure 2.**
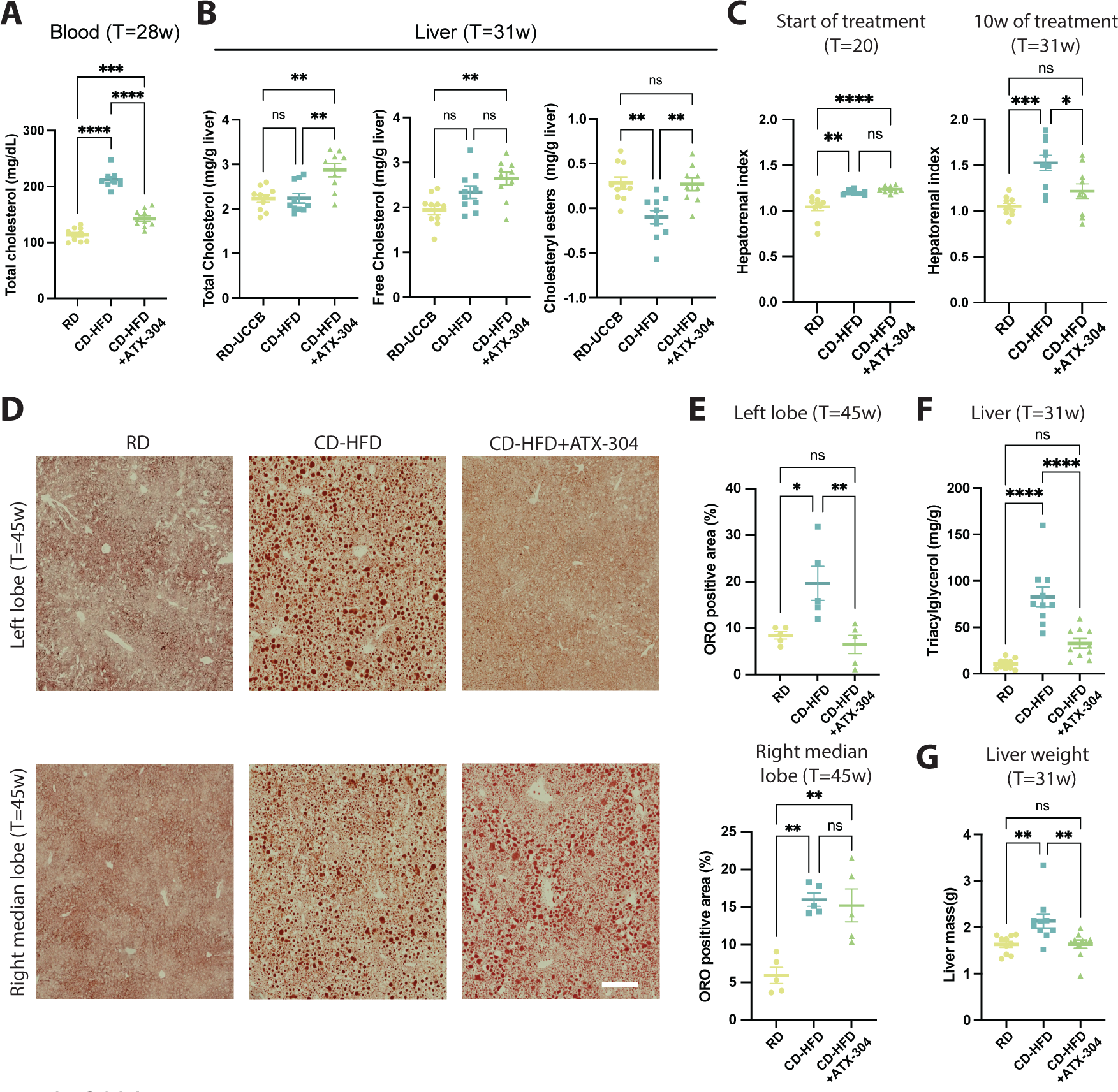
**O304 treatment decrease blood cholesterol and reduce liver lipid accumulation in obese CD-HFD mice. A**) Total blood cholesterol in RD, CD-HFD and ATX-304-treated mice at T=28w. (n=10 for all groups) **B**) Graphs depicting total cholesterol (left), free cholesterol (middle), and cholesteryl ester (right) levels in livers from RD, CD-HFD and ATX-304-treated mice at T=31w. (n=10 for all groups) **C**) Hepatorenal index of RD, CD-HFD, and CD-HFD+ATX-304 mice at start (T=21) and after 10 weeks (T=31) of treatment. (n=10 for all groups) *p<0.05, **p<0.01, ***p<0.001, ****p<0.0001 (One-way ANOVA with Tukey’s multiple comparisons test) **D**) Oil redO stained liver sections from the left lobe (upper row) and the right median lobe (bottom row) of RD, CD-HFD and ATX-304-treated mice at T=45w. Scalebar corresponds to 200um. **E**) Percentage of ORO positive area for the left (top) and right median (bottom) lobe is displayed in (n=5 for all groups). **F**) Liver trigacylglycerol content and **G**) liver mass for RD, CD-HFD and ATX-304-treated mice at T=31w (n=10 for all groups). *p<0.05, **p<0.01, ***p<0.001, ****p<0.0001 (One-way ANOVA with Tukey’s multiple comparisons test). Individual data points, mean ± SEM are indicated in all graphs.

To specifically assess the effect of ATX-304 on the livers of CD-HFD mice, animals were sacrificed after treatment and livers were collected for biochemical and histological analyses, as well as gene expression, proteomic and lipidomic profiling. Biochemical analysis revealed a ∼30% increase in total liver cholesterol in ATX-304-treated mice as compared to both healthy controls and non-treated CD-HFD mice. This increase was mainly due to an increase in the free cholesterol fraction, while the amount of inactive esterified cholesterol was restored to similar levels as control livers (Figure 2B). Given the decrease of blood cholesterol levels in treated mice, the increase in hepatic cholesterol likely reflects increased hepatic uptake. In contrast, CD-HFD livers did not present a significant increase in the amount of total or free cholesterol but were devoid of cholesteryl esters (Figure 2B). Together, these data show that ATX-304-treatment improved whole-body metabolic parameters and induced weight loss via reduced fat mass. It also shows improved blood liver profile in treated CD-HFD mice and provide evidence of an increased uptake and storage of cholesterol from the blood.

### Heterogenous reduction in liver lipid content and fibrosis in ATX-304-treated male CD-HFD mice

Ultrasound analysis before the start of treatment (T=20w) showed that the hepatorenal index (HI) was elevated in mice on CD-HFD (HI: ∼1.2) compared to RD controls (HI: ∼1.0), indicating presence of hepatic steatosis (Figure 2C). After additional 10 weeks of feeding (T=30w), the hepatorenal index was further increased (HI: ∼1.5) in non-treated CD-HFD mice. In contrast, similar to RD controls, ATX-304- treated CD-HFD mice maintained the hepatorenal index at similar levels as at the start (HI: ∼1.0 and HI:∼1.2 respectively, at T=31w). This suggested that ATX-304 treatment suppressed liver steatosis progression in CD-HFD mice. To further assess the lipid distribution in ATX-304-treated livers, Oil RedO (ORO) staining was performed on frozen sections from the left lobe and the right median lobe (Figure 2D). These analyses, together with macroscopic examination of livers during dissection, evidenced an overall reduction of lipids and lipid droplets in ATX-304-treated animals, but with substantial lobular heterogeneities (Figure 2D, Supplementary figure 3A). In CD-HFD livers treated with ATX-304, area of lipid droplets was normalized or even reduced as compared to RD controls in the left lobe for long-term treated animals, while the right median lobe maintained similar levels (Figure 2E). A similar tendency in lobular distribution was evident in short-term-treated animals (Supplementary figure 3A). Biochemical analysis also showed ∼60% reduction of triacylglycerol (TG) levels in livers treated for 10w (Figure 2F). These histological and biochemical analyses, in conjunction with the reduced total mass of the liver (Figure 2G) demonstrate that ATX-304-treatment reduced the overall lipid content of CD-HFD livers and highlights lobular heterogeneities in lipid metabolism. In the transition from simple liver steatosis to MASH, inflammatory signaling leads to activation of hepatic stellate cells and resultant fibrosis. To evaluate the level of liver fibrosis in these mice after treatment, picrosirious red (PSR) staining was applied to paraffin sections from the left lobe and the right median liver lobe (Figure 3A). The degree of fibrosis was significantly lower in the left lobe of ATX-304-treated compared to non-treated livers and normalized to that of livers from control RD mice (Figure 3B,). In contrast, fibrosis score of tissue from the right median lobe was higher in CD-HFD+ATX-304 compared with that RD and CD-HFD, showing that although there is a general amelioration of fibrosis progression, local phenotypic differences exist between the lobes (Figure 3A-B). A similar tendency was seen in the short-term cohort (Supplementary Figure 3C-D). These differences correlate with the abundance of steatosis. Taken together, these results suggest that ATX-304-treatment improves steatosis and fibrosis in male CD-HFD mice, and that these changes have a spatial pattern where the left lobe responds more strongly to the treatment regime.

**Figure 3.**
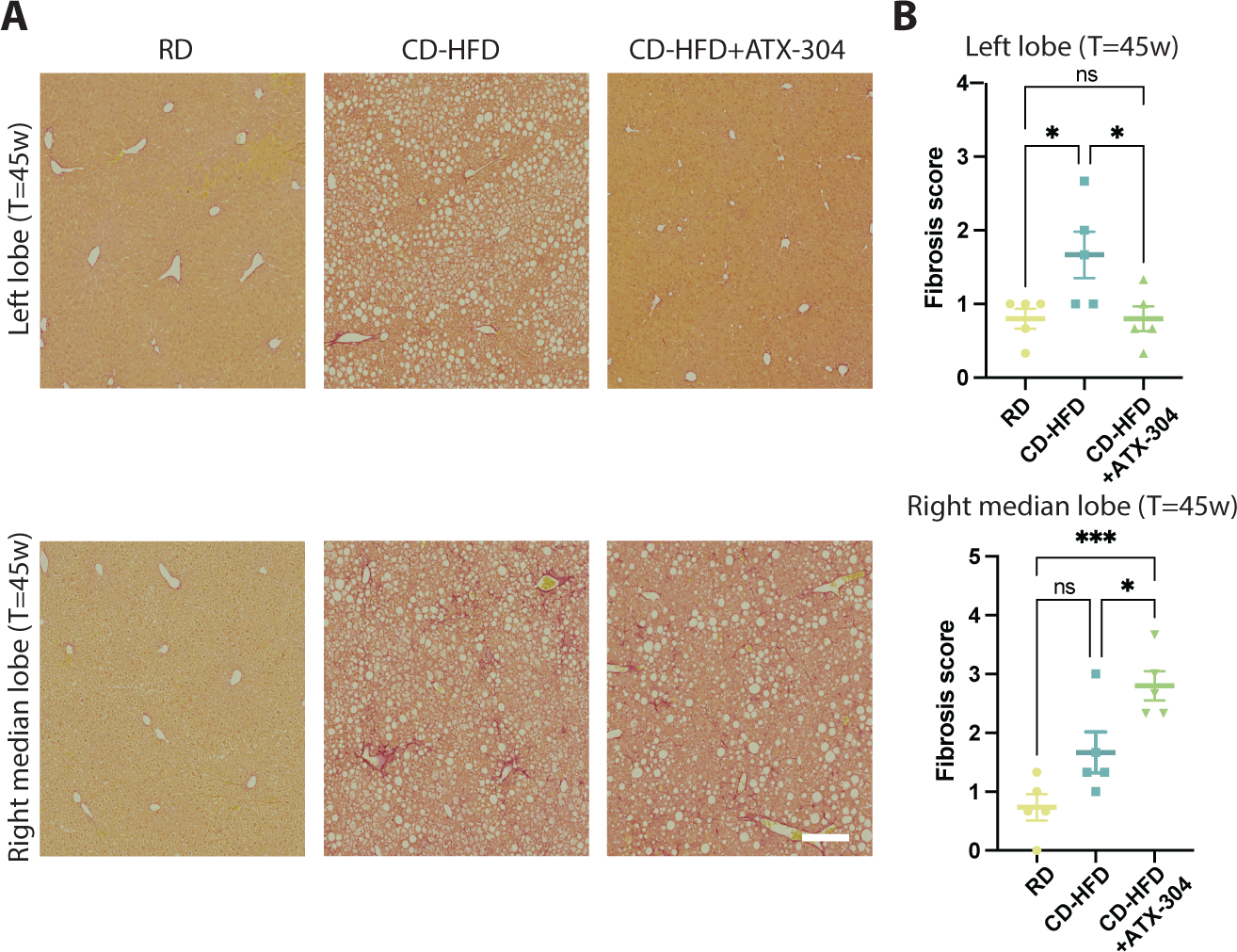
**Heterogenous amelioration of fibrosis in livers treated with ATX-304. A**) Representative images depicting picrosirius red (PSR) staining of sections from the left lobe (top row) and right median lobe (bottom row) from RD, CD-HFD and CD-HFD+ATX-304 livers. Scalebar corresponds to 200um. **B**) Fibrosis score based on PSR staining for the left (top) and right median lobe (bottom) (n=5 for all groups). *p<0.05, ***p<0.001 (One-way ANOVA with Tukey’s multiple comparisons test)

### ATX-304 mediates a transcriptional switch in liver lipid metabolism in long-term treated CD-HFD mice

To investigate the genetic underpinnings of the observed changes in lipid content and fibrosis, RNA-sequencing was performed. In livers of animals fed CD-HFD for 31w (T=31w), the number of differentially expressed genes (DEGs) compared to RD livers was 351 while for livers of CD-HFD+ATX- 304 mice (T=31w, 10w of treatment) 757 DEGS were detected (Figure 4A), Supplementary table S1). Only three genes were differentially expressed between CD-HFD and CD-HFD+ATX-304 livers (Supplementary table S1). Still, 151 DEGs were unique to CD-HFD livers and 557 DEGs were unique to ATX-304-treated CD-HFD livers (Figure 4B). Gene Ontology enrichment analysis revealed that although the most enriched GO-terms were shared between CD-HFD and CD-HFD+ATX-304 (Figure 4C), ATX- 304-treated livers were more associated to lipid and fatty acid catabolic processes and beta-oxidation (Figure 4D). Taking a closer look at key genes in lipid and cholesterol metabolism and annotating the uniquely expressed DEGs demonstrate that there is a shift in the transcriptional program of ATX-304- treated livers, suggesting a reduction in fatty acid synthesis and increased beta-oxidation, while triacylglycerol synthesis appears to be increased. Gene expression of fatty acid synthesis genes (*Acaca*, *Acab*, *Fasn)* was reduced in ATX-304-treated livers compared to RD controls, while the expression of several genes involved in beta-oxidation (*Acaa1a*, *Acaa1b*, *Acadl*, *Acadm*, *Cpt1a*, *Ehhadh*, *Hadh*, *Hadhb*) was increased (Figure 4E). The TG synthesis genes *Agpat3* and *Gpat3* were significantly upregulated in livers of ATX-304-treated mice which, given the strong reduction of fat mass in these animals, may reflect an increased influx of fatty acids from the blood (Figure 4E). This is also in line with the increased expression of genes encoding fatty acid transporter proteins *Cd36*, *Slc27a1, Slc27a2*, and the very low-density lipoprotein receptor *(Vldlr*) (Figure 4E). The latter may also contribute to an increase in the influx of cholesterol to the liver, and the reduction of blood cholesterol seen in these mice (Figure 2E). Although genes involved in cholesterol synthesis were downregulated in both CD-HFD and CD-HFD+ATX-304 (Figure 4B), this difference was more pronounced in CD-HFD livers, as illustrated by some unique DEGs for these mice (*Hmgcr*, *Mvd*, *Fdft1*, *Dhcr24*). This analysis suggests that ATX-304 mediate a catabolic switch at the transcriptional level that contribute to reduced fatty acid synthesis and increased lipid oxidation in the liver.

**Figure 4.**
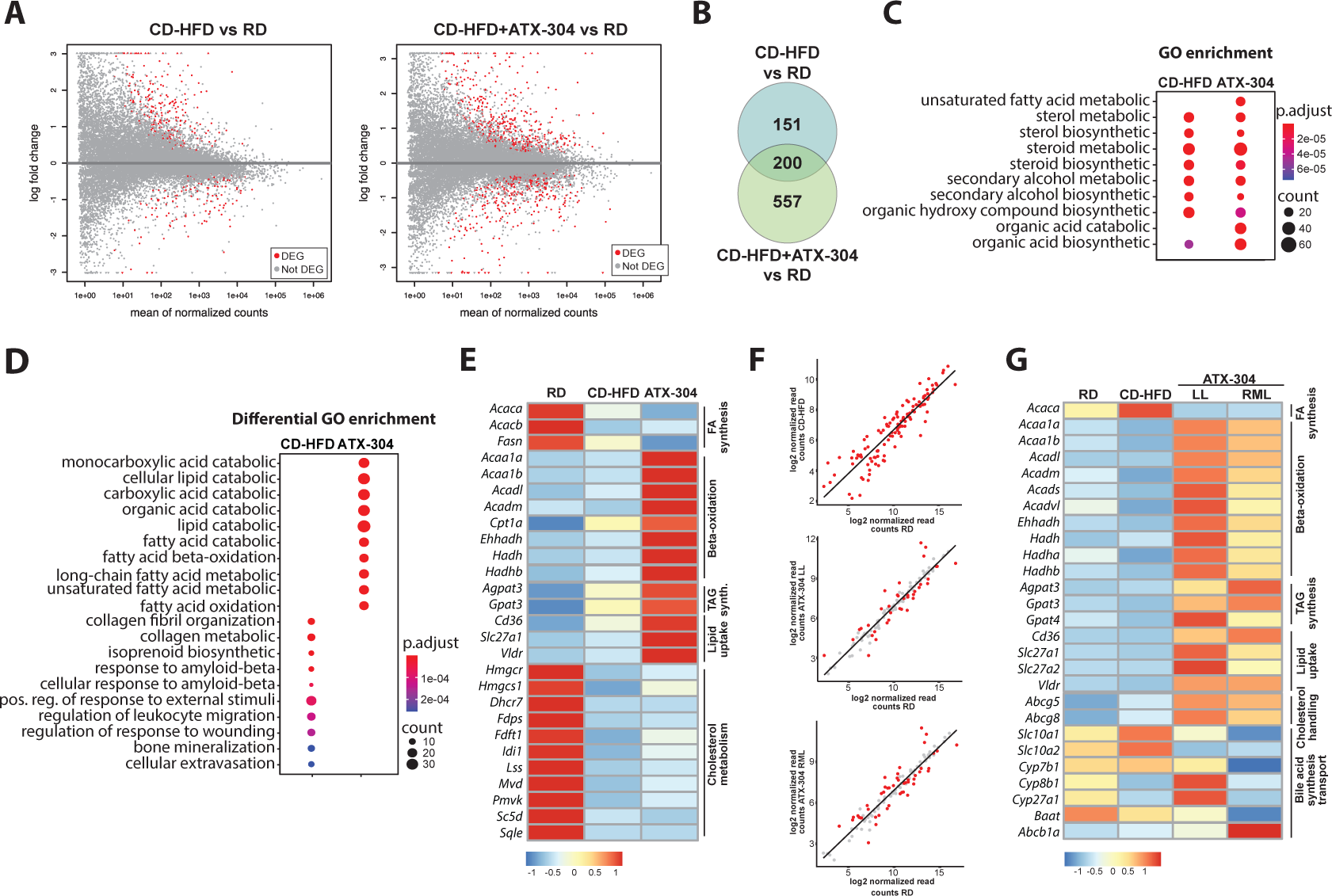
**ATX-304 mediates a transcriptional switch in liver lipid metabolism. A**) Differentially expressed genes (DEGs, red dots) in CD-HFD (left) and CD-HFD+ATX-304 (right) compared to RD control livers (*p*<0.01, FDR>0.01) **B**) Intersection of DEGs for CD-HFD and CD-HFD+ATX-304 livers. **C**) Top 10 GO terms enriched for CD-HFD and CD-HFD+ATX-304 DEGs. **D)** Bottom panel: Top 10 enriched GO terms unique for CD-HFD (left) or CD-HFD+ATX-304 DEGs (right). (p-value: hypergeometric test with Benjamini correction). **E**) Heatmap of normalized RNA-seq read counts for key lipid metabolism DEGs in long-term cohort at T=31. Scale from blue to red indicates fold change over average read count for each row. **F**) Gene expression levels of differentially expressed genes between left lobe of CD-HFD livers and RD livers (top) and the same genes plotted for the left lobe (middle) and right lobe (bottom) of HFD-ATX-304 livers compared to left lobe of RD liver. Red dots indicate genes that are differentially expressed in each condition (FDR < 0.01) **G**) Heatmap of normalized RNA-seq read counts for key lipid metabolism DEGs in livers from short-term cohort at T=10.

### Liver lipid metabolism is altered in short-term ATX-304-treated mice

Given the transcriptome diversity intrinsic to the aging process (40,41), and in view of the lobular heterogeneities in lipid distribution and fibrosis, we performed additional RNA-seq analysis for short-term treated mice (3w CD-HFD + 7w CD-HFD+ATX-304, 15-week-old). The transcriptome of the left liver lobe of RD, CD-HFD and 0304-treated CD-HFD mice, as well as the right median liver lobe from ATX-304-treated mice were analyzed. In CD-HFD livers, 115 DEGS were detected compared to RD controls. In both the left lobe and the right median lobe from ATX-304-treated livers, the expression of the majority of these genes were not significantly different from RD controls (left lobe: 78/115, right median lobe: 68/115) or changed in the opposite direction (left lobe: 6/115, right median lobe: 9/115) (Figure 4F, Supplementary table S2). As in the long-term-cohort, we detect a similar metabolic shift towards beta-oxidation and lipid transport in the treated livers, albeit more pronounced with significant differences in expression levels of key enzymes also between treated and non-treated animals. The gene expression of the fatty acid synthesis enzyme Acetyl-CoA carboxylase alpha (*Acaca*) (Figure 4G) is downregulated in both lobes of treated mice compared to CD-HFD livers and a similar trend can be seen for Acetyl-CoA carboxylase beta (*Acacb*) and Fatty acid synthase (*Fasn*) (Supplementary table S2). Several beta-oxidation genes are upregulated either in both lobes (*Acaa1a*, *Acaa1b*, *Acadl*, *Acadm*, *Acads*, *Ehhadh*, *Hadha*, *Hadhb*) or in the left lobe (*Acadvl*, *Hadh*) and the expression of TG synthesis enzymes was increased in treated animals (Both lobes: *Gpat3*, Right median lobe: *Agpat3*, *Gpat4*) (Figure 4G). We also detect significant upregulation of lipid transporters *Cd36*, *Slc27a1, Vldlr* (both lobes) and Slc27a2 (left lobe) in treated as compared to non-treated livers (Figure 4G). Interestingly, cholesterol efflux transporters *Abcg5* (both lobes) and *Abcg8* (left lobe) are upregulated in ATX-304-treated compared to RD livers, although not significantly different from non-treated livers, and a similar tendency was present in the long-term cohort. Also, bile acid transporter *Slc10a2* that is involved in cholehepatic shunting is downregulated (Figure 4G), indicating a potential increase in the excretion of cholesterol and bile. This would fit with an accommodation to higher cholesterol uptake from the blood. No major differences in metabolic programs were found between the ATX-304-treated lobes although it appears that expression of beta-oxidation genes may be slightly increased in the left lobe as compared to the right median lobe (Figure 4G). Also, genes involved in bile acid synthesis (*Cyp27a1*, *Cyp7a1*, *Cyp7b1*, *Cyp8b1*), uptake and transport (*Slc10a1, Baat, Abcb1a*) were differentially expressed in either the left or the right median lobe of the ATX-304-treated compared to non-treated livers (Figure 4G).

Taken together, these data further support the notion that ATX-304 induce a metabolic shift towards catabolism that in turn favors increased liver lipid uptake and beta-oxidation, and that also appears to increase cholesterol and bile excretion. This would thus contribute to the observed amelioration of liver steatosis, suppressed progression of fibrosis, and reduction in blood cholesterol.

### Proteomics analysis corroborate catabolic switch in lipid metabolism of ATX-304-treated livers

ATX-304 induced a clear switch in the metabolic transcriptional program of treated livers, as evidenced by RNA-seq analysis of both short-term and long-term treated mice. To investigate the effects also on the proteome, we performed proteomics analyses on mice treated with ATX-304 for 24w (T=45). We performed LC-MS/MS analysis of tissue from the left and right median liver lobe of RD, CD-HFD and ATX-304-treated mice. Given the similar proteome profiles obtained from the left and right median lobes, we considered them as whole liver and extracted a list of 138 differential proteins between CD-HFD and ATX-304-treated livers (Supplementary table S3). GO analysis of differential proteins revealed a significant enrichment for terms related to fatty acid metabolism (Supplementary table S4). In CD-HFD+ATX-304 livers compared to CD-HFD livers, abundancy of proteins involved in activities such as ’fatty acid hydroxylase activity,’ ’fatty acid hydrolase activity,’ ’fatty acid monooxygenase activity,’ and ’steroid dehydrogenase activity’ were significantly increased while proteins involved in ’carboxylase and desaturase activity of co-enzymes involved in fatty acid metabolism’ and ’fatty acid acyl transferase related synthase activity’ were significantly decreased. Similarly, KEGG (Kyoto Encyclopedia of Genes and Genomes) pathway enrichment analyses identified pathways linked to fatty acid degradation and biosynthesis, PPAR-signaling, as well as metabolism (Figure 5A).

**Figure 5.**
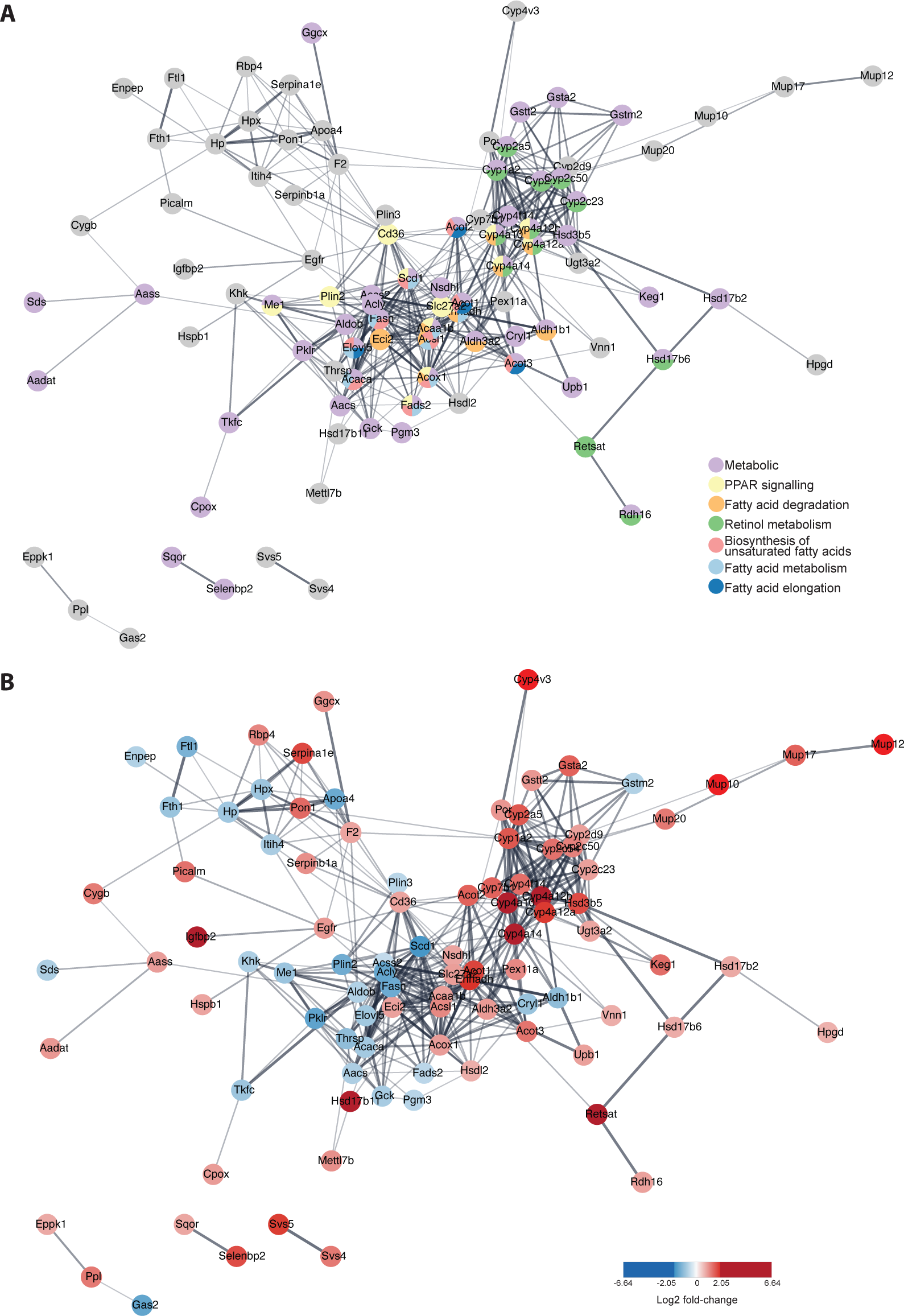
Proteomics analysis corroborate catabolic switch in lipid metabolism of ATX-304-treated livers. STRING protein interaction network of differential proteins depicting enriched KEGG pathways in **A**) and log2 fold-change of individual proteins in **B**).

Key enzymes involved in beta-oxidation (ACAA1B, EHHADH, ACOX1) and lipid transport (CD36, SLC27A2) were significantly increased in treated livers compared to both RD and CD-HFD livers, while proteins involved in fatty acid activation (ACSL1, ACSL4) were increased in comparison to CD-HFD- livers only (Figure 5B, Supplementary table S3). On the contrary, fatty acid synthesis enzymes ACACA and FASN were more abundant in CD-HFD compared to RD livers with no difference between ATX- 304-treated livers and RD controls (Figure 5B, Supplementary table S3). Taken together, this further demonstrate that ATX-304 mediate a catabolic switch that leads to reduced fatty acid synthesis and increased lipid oxidation in the liver. This aids in suppressing lipid accumulation in treated livers, despite the potential increased lipid influx as suggested also by the increased levels of lipid transporters.

### ATX-304-treatment induce remodeling of lipid zonation and ameliorate lipid profile alterations in CD-HFD livers

Histological analysis of lipid droplet distribution did not only reveal reduced lipid content and lobular differences in treated livers, but also demonstrated a switch in lobular zonation pattern. As expected, both RD control livers and CD-HFD livers presented lipid droplets predominantly in the pericentral zone of the liver lobules (Figure 6A). In contrast, ATX-304-treated animals consistently presented a higher proportion of lipid droplets towards the periportal region (Figure 6A).

**Figure 6.**
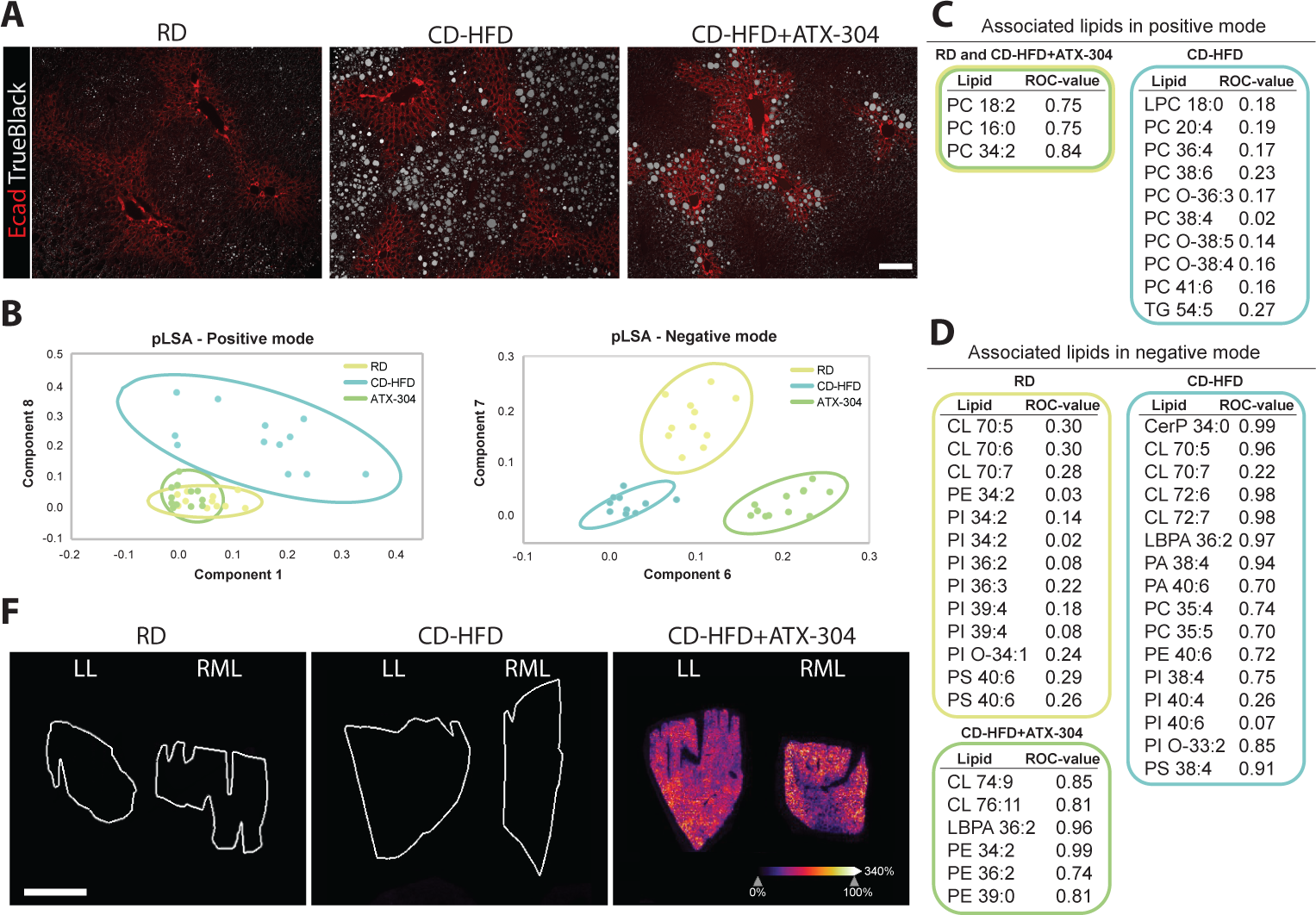
**ATX-304-treatment ameliorate lipid profile alterations in CD-HFD livers. A**) Overlay of immuno- flourescence and brightfield images of liver sections stained for periportal marker E-cadherin (red) and with TrueBlack for lipid droplets (white). Lipid droplets in ATX-304-treated livers are preferentially located in the periportal zone. Scale bar corresponds to 50 um. **B**) pLSA score plot for all MALDI-MSI samples in positive mode. **C**) Lipid species identified in positive mode and associated with RD and CD-HFD+ATX-304 livers (yellow and green box), or CD-HFD livers (blue box). ROC-values are indicated. **D**) pLSA score plot of all MALDI-MSI samples in negative mode. **E**) Lipid species identified in negative mode and associated with RD (yellow box), CD-HFD (blue box) and CD-HFD+ATX-304 (green box). **F**) Images of O304 localisation at 378.012m/z ± 0.3Da in the left (LL) and right median liver (RML) lobe of treated animals. RD and CD-HFD livers display no signal. White scale bar corresponds to 2mm. Graded scale from blue (0%) to white (100%) indicate signal intensity.

It is well known that lipogenic processes such as fatty acid and triacylglycerol synthesis are more active pericentrally while beta-oxidation primarily occur periportally (42). In addition, recent mass spectrometry imaging has shown zonal distribution of specific lipid species and alterations in these patterns upon MASDL and MASH (Hall.2017.Hepatology, Seubnooch.2023.JHEP Reports). To further investigate the effects of ATX-304-treatment on the lipid distribution in CD-HFD livers, we performed matrix-assisted laser desorption/ionisation mass spectrometry imaging (MALDI-MSI) on sections from the left and right median liver lobes of all three experimental groups. Using this method, we were able to identify lipids of interest while preserving the spatial distribution. Given the absence of pronounced differences between the lobes, the lipid profile of each liver condition was generated by merging this data. We identified several lipid groups such as lysophosphatidylcholines (LPC), oxidized phosphatidylcholines (oxPC), and TGs in positive mode, while Ceramides (Cer), phosphatidylethanolamines (PE), phosphatidic acid (PA), phosphatidylinositols (PI), phospatidylcholines (PC), phosphatidylserines (PS), Lysobisphosphatidic acid (LBPA), and cardiolipins (CL) were found in negative mode.

Interestingly, unsupervised pLSA analysis of the positive ion mode data demonstrated that the CD-HFD livers could be clearly separated from RD controls and ATX-304-treated livers (Figure 6B). These findings indicate that the lipid profiles of the ATX-304-treated livers are more similar to RD controls and thus appear to have been partially restored. In addition, several species of oxidized lipids were characteristic of the CD-HFD samples (Figure 6C), indicating that oxidative stress may be higher in these livers compared to both ATX-304-treated livers and RD controls. Additional differences between the groups were detected in the abundance of distinct PC species, while TG 54:5 and LPC 18:0 were also more abundant in CD-HFD livers (Figure 6C). Of note, high levels of LPCs have been associated with disrupted mitochondrial integrity, inflammation, and apoptosis (44).

In negative mode, pLSA analysis separated all three groups in different clusters, indicating also distinct lipid profile features for each of the groups (Figure 6B). Among the differential lipids, 15 were distinctive for CD-HFD, 12 for RD, and 6 for ATX-304-treated livers (Figure 5E). The most abundant among differential lipids were CL (9) and PI (11), but also LBPA (2), PA (2), PC (2), PE (3), and PS (3) had significantly different abundance. All differential lipids obtained in positive or negative mode were further validated by ROC analyses (Figure 6C and D). All the above-mentioned lipids were homogenously distributed across the liver lobes and did not correlate with the observed switch in zonal distribution of lipid droplets.

To further investigate the potential mechanism underlying changes in lipid droplet distribution, e. g. via differential metabolic activity due to heterogenous uptake of ATX-304, MALDI-MSI was also employed to visualize the spatial distribution of ATX-304 in both the left and right lobes of the dosed animals (Figure 6F). These analyses revealed that the drug was evenly distributed in the livers of dosed animals, with similar high intensities between the two lobes (Figure 6F). As expected, no signal was found in RD and CD-HFD livers (Figure 6F). MS/MS was used to confirm the identification of the drug (Supplementary figure 4). Thus, changes in lipid zonation are not due to variation in ATX-304 uptake, nor is the lobular heterogeneities in lipid content.

Taken together, these data demonstrate that ATX-304-treatment of CD-HFD mice leads to both quantitative and qualitative changes in liver lipid abundance, including a reduction in oxidized lipids suggestive of an amelioration of MASDL, and a shift in lipid profile towards a more normal composition.

## Discussion

The idea that AMPK holds promise as a potential target for treatment in MASLD and MASH has been around for many years. Despite efforts to test various AMPK activators in animal models, no direct AMPK activator has yet reached the clinic as treatment for this metabolic disease. Here, we investigated the effects of the clinical stage AMPK-activator ATX-304 (28) in a CD-HFD mouse model of MASLD.

Our data show that ATX-304-treatment reduces whole-body fat mass, reduces hepatic steatosis, and suppresses liver fibrosis in mice fed CD-HFD. It is well documented that AMPK activation inhibits *de novo* lipid synthesis (13,15,18,24,45) and accumulating evidence also suggest simultaneous stimulation of fatty acid oxidation (13,16,18). In ATX-304-treated mice we detected changes at the level of both the gene and protein expression, in accordance with a metabolic shift in the liver favoring beta-oxidation at the expense of fatty acid synthesis. In white adipose tissue, AMPK is known to inhibit lipogenesis while the reports of effects on lipolysis are more contradictive (16,25,46–49). Notably, the idea that hepatic activation of AMPK trigger mobilization of fat from the adipose tissue (16) fits with several of our observations. Firstly, the almost complete reduction of whole-body fat mass suggests that ATX-304 mediates a metabolic shift towards lipid catabolism, which can also lead to an increased release of fatty acids into the circulation. Secondly, we detected upregulation of fatty acid transporters in the livers of treated animals, and zonation of lipid droplets are reversed in these mice. Alike beta-oxidation, evidence suggests that fatty acid trafficking occur preferentially in the periportal zone of the liver lobules. It has been shown that there is a higher blood concentration of fatty acids in the periportal region, fatty acid binding proteins are more highly expressed (50,51), and that glucagon induce higher uptake of fatty acids in periportal than pericentral hepatocytes (52). Although the general lipid content is reduced in livers of ATX-304-treated animals, our data suggest that in some regions with more lipids, increased periportal beta-oxidation does not suffice to utilize all the fatty acids taken up from the circulation and these are used for TG synthesis. In contrast, pericentrally, lipid influx may be lower and the combination of reduced lipogenesis and increased beta oxidation would be sufficient to reduce the abundance of lipid droplets. This balance between influx, beta oxidation and lipid synthesis could potentially explain both the local lobular heterogeneity in lipid abundance, but also the reversal of the lipid droplet zonation pattern seen in livers of ATX-304-treated mice. Along these lines, short-term over-activation of AMPK in the liver also leads to a concomitant initial accumulation of lipids in the liver and increased beta-oxidation (53), which is also observed in the physiological response to fasting (54). Thus, these observations underline the importance of the fine-tuning between lipid utilization, lipid synthesis and fatty acid uptake to determine phenotypic output in terms of cellular lipid content and in the long term its impact on liver function.

In line with previous reports that pharmacological AMPK-activation reduce blood cholesterol in both rodents and primates (18), we see a significant reduction of blood cholesterol in CD-HFD mice treated with ATX-304. Based on gene expression data and the fact that hepatic cholesterol levels are increased in these mice, it is likely that this reduction in blood cholesterol is due to increased hepatic uptake rather than reduced cholesterol synthesis in this model. Total hepatic cholesterol is more abundant in ATX-304-treated livers compared to both CD-HFD livers and controls but importantly, this increase includes a normalization of esterified cholesterol levels and an increase in the ratio of esterified cholesterol to free cholesterol compared to CD-HFD. This is indicative of an improvement in the cholesterol handling capacity as MASLD is associated with accumulation of free cholesterol without concomitant increase of esterified cholesterol (55). The increase in blood bile acids is likely also connected to changes in the metabolic pathways of cholesterol.

Apart from reduced hepatic lipid content, ATX-304-treatment induced qualitative differences in lipid species composition compared to CD-HFD, even to some extent restoring the lipid profile of RD control livers. More importantly, lipid profiling revealed that oxidized lipids were more abundant in livers from CD-HFD animals compared to both RD controls and ATX-304-treated livers. Oxidized lipids are formed when reactive oxygen species (ROS) react with various lipid species, and increased levels of oxidized lipids are associated with the progression of simple steatosis to MASH (56–58). Our data suggest that ATX-304-mediated activation of AMPK reduces the production of oxidized lipids in the liver thus preventing the buildup of these harmful lipid species. This is also in agreement with other studies showing that AMPK activation mediates responses to ameliorate oxidative stress and suppresses ROS formation via induction of antioxidants and mitochondrial uncoupling (59–61) . Interestingly, it was recently demonstrated that ATX-304 induces mitochondrial uncoupling in C2C12 myotubes (31). Similar mechanisms may be in place in ATX-304-treated livers and would provide an attractive hypothesis to at least partially explain how the hepatocytes cope with the increased lipid load upon activation of catabolic programs.

In conclusion, this study emphasizes the therapeutic potential of ATX-304 in fatty liver disease including decreased hepatic lipid content and body adiposity, suppression of fibrosis, and lowering of blood cholesterol, and at the same time highlight lobular heterogeneities in lipid metabolism in the murine liver.

## Experimental procedures

### Animals and diets

All experiments were performed in compliance with national and institutional laws and are reported in accordance with the ARRIVE guidelines. The study was approved by the Regional Ethics Committee at the Court of Appeal of Northern Norrland (Ethical approval ID A39-2018). Our study examined male mice as they have been reported to have more severe steatosis and fibrosis in this model. C57Bl6 mice were purchased from Charles River Lab and housed at 12:12 h light/dark cycle in a temperature/humidity controlled (22°C and 50% humidity) room with *ad libitum* feeding. 5-week old male C57Bl6 mice were fed with either standard chow (RD, Special Diet Service #801730) or choline deficient high-fat diet (CD-HFD, *D05010402,* Research diets). After 3 weeks (3w, short-term cohort) or 21 weeks (21w, long-term cohorts) half of the mice fed CD-HFD were switched to CD-HFD supplemented with 1 mg/g ATX-304 (CAS # 1261289-04-6, kindly provided by Betagenon AB, Umeå, Sweden). Mice in the short-term cohort were sacrificed 7weeks (7w) after diet-switch while long-term cohorts were sacrificed after 10 or 24 weeks (10w or 24w) of treatment. Food intake and body weight were measured weekly.

### Body composition, blood profiling and measurement of hepatorenal index

Body composition (fat and lean weight) of live mice was measured using EchoMRI (EchoMRI LLC, EchoMRI 3-in-1). Blood profiling was performed on 100 µl of freshly collected blood using the Mammalian Liver Profile rotor on a VetScan VS2 (Abaxis, Inc.). Blood glucose levels were measured using Glucometer (Ultra 2, One Touch) and plasma insulin was analysed via the ultrasensitive mouse insulin ELISA kit (Chrystal Chem Inc. #90080). Hepatorenal Index was measured using the Vevo 2100 system and the MS550D transducer (Fujifilm VisualSonics, Toronto, ON, Canada). Hepato-renal index was calculated as the intensity in kidney cortex echogeneity, divided by the liver intensity. Three different images per mouse were used to calculate a mean.

### Liver triacylglycerol and cholesterol measurements

Liver triacylglycerol (TG) was measured using Serum Triglyceride Determination Kit (Sigma-Aldrich #TR0100) according to manufacturer’s recommendations. Briefly, 0.2-0.3 g of liver were homogenized in 2 ml PBS before addition of 6 ml chloroform/methanol (2:1). Samples were mixed until phase separation no longer occurred and left at RT 30 min before centrifuged at 4500 rpm for 5 min. The chloroform phase was transferred into pre-weighted glassware and kept at 4°C O/N. Any water drops were removed and the chloroform evaporated by a stream of nitrogen before residual solvent was removed in a SpeedVac, 15 min. The glassware was re-weighed, and total lipids were calculated (mg/g liver). The residue was dissolved in 35% Triton X-100/methanol. Liver triglycerides were determined with the Serum Triglyceride Determination Kit (Sigma-Aldrich #TR0100) according to the manufacturer’s recommendations with a minor modification for triglyceride determination, which was analysed at 560 nm instead of 540 nm.

Liver cholesterol was measured using the Cholesterol / Cholesteryl Ester Quantitation Kit (MBL #JM*-* K603*-*100) according to manufacturer’s recommendations. Briefly, 25 mg of liver was homogenized in PBS. Chloroform, isopropanol and NP-40 (7:11:0.1) were added, followed by vortexing and centrifugation at 15000 *g* for 5 min. The water phase was removed, and chloroform evaporated with nitrogen gas. The sample was dissolved and diluted in cholesterol assay buffer and loaded onto a 96- well plate with reaction mix (Cholesterol reaction buffer, probe, enzyme mix, and cholesterol Esterase). The plate was incubated in dark for 1 hour at 37 °C. Absorbance measurement was done at 570 nm using an ELISA plate reader.

### Liver histology, fibrosis scoring and lipid droplet distribution

Liver histology and fibrosis were assessed by haematoxylin & eosin (H&E) and picrosirius red staining on paraffin-embedded formalin-fixed sections. Paraffin-embedded liver tissues were sectioned and stepwise deparaffinized in ethanol (100%, 95%, 70%, 50%), rinsed with water. Mayers HTX PLUS (HistoLab, #01825) and eosin (51500 mg Eosin Y in 79% ethanol with 25*10^-5^%acetic acid) were applied sequentially for 2 min followed by a 5 min wash in running water in between. Slides were then rinsed in xylene and mounted with DPX mounting media (Sigma-Aldrich, #1.00579.0500) before image acquisition.

Picrosirius red was applied for 1 hour followed by two washes with acidified water (0.5 % glacial acetic acid in MQ). Excess water was removed, and sections were dehydrated in ethanol (2 x 100% EtOH) followed by clearing in xylene and mounting with DPX mounting media (Sigma-Aldrich, #1.00579.0500). Fibrosis was assessed using the Ishak fibrosis scoring system (33). The mean score of three sections was used for each mouse (n=5 for all experimental groups).

Lipid droplet distribution was assessed by Oil Red O (ORO) staining on cryo-embedded liver tissue (Frozen in OCT after 4% PFA fixation). Tissues were sectioned, dried at RT for 10 min and washed in PBS followed by a quick rinse in 60% isopropanol. Sections were stained with ORO working solution for 15 min, rinsed in 60% isopropanol, and washed two times for 5 min in MQ before mounting with Vectashield mounting media (Vector Laboratories, #H-1000). Lipid content was quantified using Qupath version 0.2.2. For each sample, the mean of three quantified sections was used (n=5 for each experimental group). All slides were imaged on an Axioscan Z1 slidescanner (ZEISS).

### RNA-seq

#### Library preparation

Approximately 30 mg of frozen liver were used for total RNA extraction using the RNeasy Plus Universal minikit (Qiagen, #73404). Integrity and quality of RNA was determined on a Bioanalyzer (Agilent, #5067-1511). Poly(A) RNA was purified with the NEBNext Poly(A) mRNA Magnetic Isolation Module (NEB, E7490) and RNA-seq libraries were prepared using the NEBNext Ultra II RNA Library Prep Kit from Illumina (NEB, E7770) following the manufacturer’s instructions. The quality and concentration of libraries were assessed with Bioanalyzer (Agilent, #5067-4626) and Qubit (Thermofisher, #Q33230). Libraries were sequenced on a NovaSeq 6000 Illumina Sequencer, with a coverage of ∼ 37 million 150PE reads per library, and quality control of fastq files was done with FastQC.

#### Gene expression analysis

Raw reads were aligned to the mouse genome (mm10, UCSC) using STAR (options: -- outSAMtype BAM SortedByCoordinate --seedSearchStartLmax 12 -- outFilterScoreMinOverLread 0.3 -- alignSJoverhangMin 15 --outFilterMismatchNmax 33 -- outFilterMatchNminOverLread 0 -- outFilterType BySJout --outSAMattributes NH HI AS NM MD --outSAMstrandField intronMotif -- quantMode GeneCounts) (34). Genes with a minimum of 10 reads along all the samples were kept for further analysis. Normalisation and differential expression analysis were performed using DESeq2 (35). Genes were considered differentially expressed with p-value < 0.01 and FDR < 0.01 (False Discovery Rate). Gene Ontology (GO) and KEGG enrichment analysis for each cluster was performed using clusterProfiler (pvalue < 0.01 and FDR < 0.01) (36).

### Proteomics

#### Chemicals and reagents

Ammoniumbicarbonate (ABC), dithiothreitol (DTT), iodoacetamide (IAM), formic acid (FA, ULC grade) and xylene were purchased from Sigma-Aldrich (Zwijndrecht, The Netherlands). Rapigest SF was obtained from Waters (Milford, USA). Trypsin/Lys C was purchased from Promega (Madison, USA). ITO slides were obtained from Delta technologies (Loveland, USA). Solvents including methanol, chloroform were purchased from Biosolve Chimie SARL (Dieuze, France). For analysis of drug distribution, 1 mg of the ATX-304 (Betagenon) was dissolved in 1 mL methanol (final concentration of 1mg/mL)

#### Sample preparation for label free proteomics

A piece of each liver lobe was dissected and placed in 100 µL urea buffer (5 M). Tissue was homogenized 10 seconds in a mini-bead beater using 1.0 mm glass beads (BioSpec Products, Bartlesville, OK, USA) at 2500 rpm. To lyse the cells, three freeze-thaw cycles were performed. Bradford assays were performed to determine protein concentration, and samples were diluted in 50 µL urea buffer (50 mM ABC / 5 M Urea) with a protein yield of 37.5 µg per sample. For protein extraction, 5 µL of DTT solution (200 mM) were added, followed by vortexing and incubated 45min at 21°C. Then, 6 µL of IAM (400 mM) was added, samples were vortexed, and then incubated 45min at 21°C in darkness. 10 µL of DTT solution (200 mM) was added to the samples, followed by vortexing and incubation 45min at 21°C. Samples were further incubated 2h at 37°C with Trypsin/LysC in a 1:25 (1.5 µL) enzyme to protein ratio, followed by overnight incubation at 37°C in a thermomixer (250 rpm) upon addition of 200 µL of ABC (50mM). The tryptic digestion reaction was stopped by addition of 30 µL of 20% ACN / 10% FA. To eliminate any remaining particles, the samples were centrifuged for 30 min (15,000 g, 4°C), supernatants were collected and stored at -80°C until further processing.

#### LC-MS/MS label free proteomics

Separation of peptides was performed on a Thermo Scientific (Dionex) Ultimate 3000 Rapid Separation UHPLC system equipped with a PepSep C18 analytical column (15 cm, ID 75 μm, 1.9 μm Reprosil, 120 Å) coupled to a Q Exactive HF mass spectrometer (Thermo Scientific). An aliquot of 2 μL (tissue piece) or 10 μL (tissue section) was injected and desalting was performed using an on-line installed C18 trapping column. After desalting, peptides were separated using a 90 min linear gradient going from 5 to 35% ACN with 0.1 % FA at 300 nL/min flow rate. Mass spectra were acquired between m/z 250- 1250 with a resolution of 120.000 in positive polarity. This was followed by MS/MS scans in DDA mode of the top 15 most intense peaks with a resolution of 15.000.

#### Data processing proteins

Proteome Discoverer software version 2.5 (Thermo Scientific) was used to identify proteins by processing raw files with the search engine Sequest and the databases Homo Sapiens (TaxID 9606, version 2020-03-25) and Mus musculus (TaxID 10090, version 2022-03-15). The tolerance for precursor mass was set at 10 ppm, with a fragment tolerance of 0.02 Da. Trypsin was selected as an enzyme with no more than two missed cleavage sites. The total peptide count was used to normalize the data. The FDR was set to a maximum of 1%. Statistical significance of protein abundance differences between CD-HFD+ATX-304 and CD-HFD livers was assessed by a one-way ANOVA test. Proteins were considered significantly different between two conditions if the adjusted p-value was less than 0.05 and the fold-change (FC) threshold was ± 1.5.

#### Protein pathway analysis

To identify the top enriched pathways, all modulated proteins were loaded into the STRING database program (string-db.org, version 11.5) and EnrichR software. These pathways were considered relevant if the adjusted p-value was less than 0.05, and they were then ordered based on their combined score in EnrichR. For pathway analysis, the Kyoto Encyclopedia of Genes and Genomes (KEGG) database was used. The pathways were visualized using the STRING database and ordered based on the calculated strength in STRING database. This strength is calculated by the ratio of the number of proteins in our network that are annotated and the expected numbers of proteins in a random network of the same size.

### Matrix-Assisted Laser Desorption/Ionisation (MALDI) - Mass Spectrometry Imaging (MSI)

#### Chemicals and reagents

Norharmane was purchased from Sigma-Aldrich (Zwijndrecht, The Netherlands). ITO slides were obtained from Delta technologies (Loveland, USA). Solvents including methanol, chloroform were purchased from Biosolve Chimie SARL (Dieuze, France). For analysis of drug distribution, 1 mg of ATX- 304 (Betagenon) was dissolved in 1 mL methanol (final concentration of 1mg/mL)

#### Matrix Application

Norharmane (Sigma-Aldrich) was utilized for lipid analysis. The HTX TM-Sprayer system (HTX technologies, LLC, NC, USA) was used to spray 15 layers of the matrix at a concentration of 7 mg/mL in chloroform/methanol (2:1, v/v) at a flow rate of 0.12 mL/min. The N2 flow rate was set at 10 psi. A drying period of 30 seconds was employed between each layer, and a velocity of 1200 mm/min was used.

#### Data acquisition of lipids and their fragments

For each of the three conditions (n=3, 3 animals per condition) MALDI-MSI experiments were carried out in both polarities (positive and negative) utilizing a RapifleX MALDI Tissue-typer instrument in reflectron mode (Bruker, Bremen, Germany). Spectra were acquired in the mass range *m/z* 300-1600. The molecular profiles of all conditions were compared using a spatial resolution of 30 μm. After analyzing a sample in positive mode, the same section was re-analyzed in negative mode with an offset of 15 µm in the x-direction to avoid sampling the same location.

To structurally identify lipids, additional MALDI-MSI MS/MS experiments were performed on consecutive sections. This was done on an Orbitrap-Elite hybrid ion trap MS instrument (Thermo Fisher Scientific GmbH, Bremen, Germany) running in data-dependent acquisition (DDA) mode^1^. MS1 data was acquired over a mass range of *m/z* 400-1600 with a resolution of 240.000 @ 400 *m/z*. MS2 data was acquired in the ion trap with an isolation width of 0.7 *m/z*. In positive mode, a collision energy of 30 eV was applied, while in negative mode, a collision energy of 38 eV was used. These collision energies were fixed for the whole mass range. The stage step size was set at 25 μm (horizontal) x 50 μm (vertical) and measured in positive and negative polarity.

#### Identification of ATX-304

MS/MS of ATX-304 was performed on a timsTOF flex (Bruker Daltonics Inc.) in negative polarity. The isolation window was set at 378.00 ± 1.00 m/z. Collision energies of 20 eV and 50 eV were used. The method was used on a spotted drug standard (1 µL spotted on an ITO slide) as well as on a dosed liver section. Both were sprayed with norharmane using the same settings as mentioned before.

#### Lipid data analysis

To compare the lipid profiles between RFD, CD-HFD and CD-HFD+ATX-304, MALDI-MSI data was imported and analyzed in SCiLS lab software (SCiLS, Bremen, Germany). From SCiLS, the overview spectra were exported to cvs format and imported into mmass (5.5.0, www.mmass.org). Afterwards, peak picking was performed in Mmass with the following settings for both polarities: signal-to-noise of 3, relative intensity threshold of 0.5 %, peak height of 50%. Baseline smoothing was applied with precision of 11 and an offset of 41. To exclude matrix clusters, a mass range of *m/z* 500-1600 was used. After peak picking, the spectra were imported back into SCiLS and normalized by total ion current (TIC). Probabilistic latent semantic analysis (pLSA) with random initialization was performed on the data with an *m/z* interval set to 0.3 Da for all comparisons, using the mean spectra (mean spectrum of each section separately). To validate the *m/z* values found in the pLSA, additional Receiver Operating Characteristic (ROC) analysis was performed with a cut-off value above 0.70 or below 0.30 for the area under the ROC curve (AUC) to assign peaks as discriminative. To identify lipids species with zonal distribution, pLSA was performed on all samples by using the individual spectra (spectrum of each pixel) on the data normalized by TIC (*m/z* interval 0.3 Da) and compared to histology and immunostaining from consecutive sections. The mass range for this analysis was set to *m/z* 300-1600.

#### Data processing of lipid fragments for lipid Identification

The parent ion masses obtained from the raw full scan FTMS were used for lipid identification with their corresponding generated IT-MS/MS spectra. All MS/MS files were converted into imzML and imported into LipostarMSI (37)for further analysis. The Lipid Maps database (edition July 2020) was used for lipid identification, such that the [M-H]^-^ ion or [M-H]^+^ ion was selected at a mass tolerance of 0.3 Da ± 5 ppm. MS/MS analyses are performed on the precursor [M-H]^-^ ion or [M-H]^+^ ion and fragments were matched with a m/z tolerance of 0.25 Da ± 5ppm. Only lipids that have a minimum carbon chain length of 12 used for identification in LipostarMSI.

Identification of the cardiolipins was done manually by comparing m/z values from our high mass resolution orbitrap data with m/z values published in the literature (38,39). In addition, at least two fragments of each cardiolipin were manually annotated using the raw data. Supplementary S5 contains a table with the ppm errors of all identified lipids.

## Author contributions

The order of the first two authors was determined alphabetically. E.H. acquired, analysed and interpreted the data, and contributed to writing the original draft. I.V. performed lipidomics and proteomics data acquisition and analysis, and contributed to writing the original draft. S.P. contributed to tissue collection, acquisition and analysis of histological and biochemical data. A.L.-P. generated sequencing libraries and performed bioinformatics analysis. B.C.P. and M.V. supervised lipidomics and proteomics data acquisition and analysis. S.R. contributed to conceptualisation and experimental design, and co-supervised the bioinformatics analysis. A.H. conceptualised and directed the research, secured funding, analysed and interpreted the data, and wrote the final manuscript with input from E.H., I.V., B.C.P., M.V. and S.R.

## Acknowledgements

We thank Ingela Lundberg for animal care and Madeleine Ericsson for technical assistance with measurements of hepatorenal index. We also acknowledge the facilities and technical assistance of Umeå Center for Comparative Biology (UCCB). ATX-304 was kindly provided by Balticgruppen Bio AB/Betagenon, Umeå, Sweden. The computations were enabled by resources provided by the National Academic Infrastructure for Supercomputing in Sweden (NAISS) and the Swedish National Infrastructure for Computing (SNIC) at UPPMAX partially funded by the Swedish Research Council through grant agreements no. 2022-06725 and no. 2018-05973. The research leading to these results has received funding from the Kempe Foundation (SMK-1863 & JCK-2149), The Cancer Research Foundation Norrland (Grant Nos. AMP 18-940 and AMP 21-1043), and Lion’s Cancer Research Foundation in Northern Sweden (LP 20-2232 and LP 22-2313). We also acknowledge support from the Strategic Research Program in Diabetes at Umeå University. Finally, we acknowledge past and present members of the Hörnblad and Remeseiro laboratories for helpful discussions.

