## Supplementary_material for "AMPK-activator ATX-304 reduces oxidative stress and improves MASLD via metabolic switching"

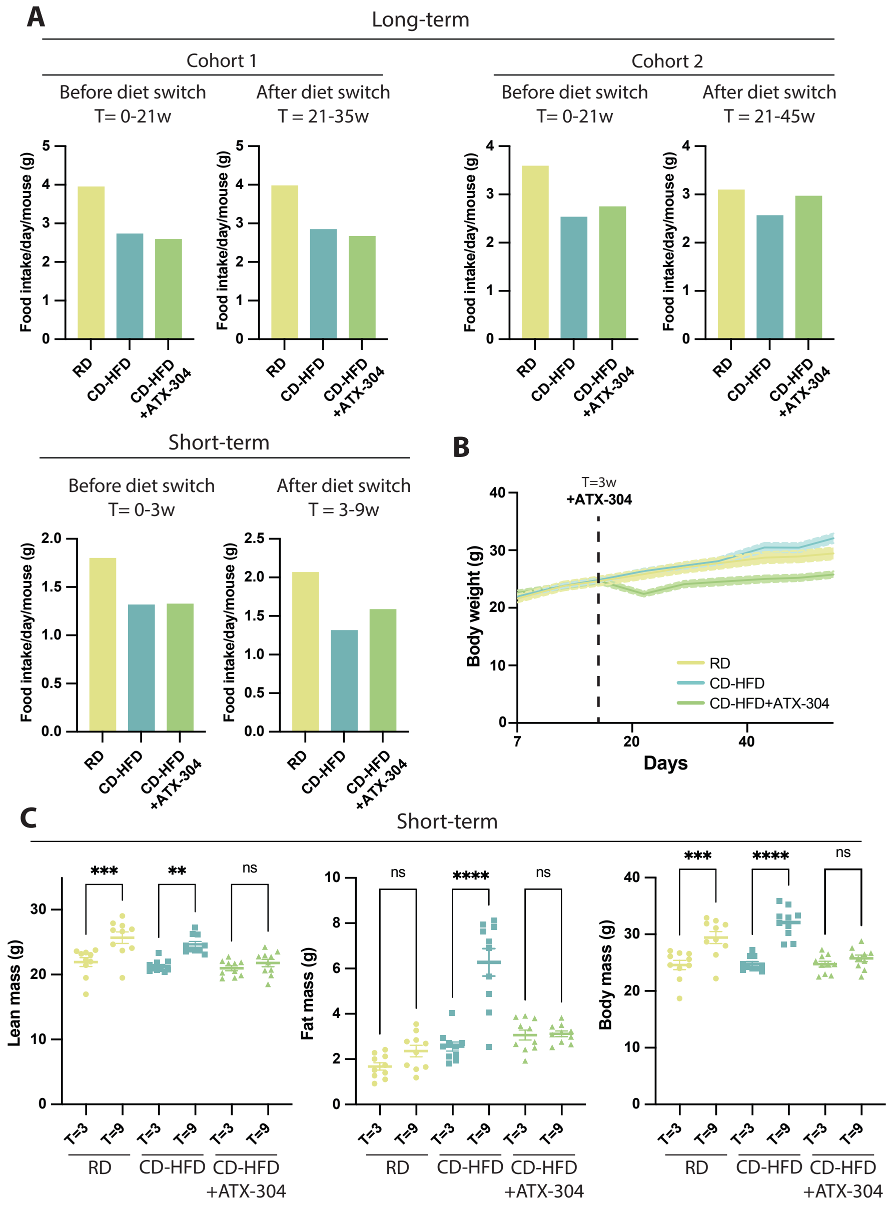


**Supplementary Figure 1. ATX-304 does not induce decreased food intake in C57Bl/6J male mice. A**) Average food intake per day and mouse for RD, CD-HFD and CD-HFD+ATX-304 in long-term (top row) and short-term cohorts (bottom row). **B**) Weight curve for short-term RD, CD-HFD and CD-HFD+ATX-304 mice. Dashed line indicate start of ATX-304 treatment (T=3w). Colored shade depicts standard error of the mean (SEM). **C**) EchoMRI measuring fat, lean and total mass(g), before and after ATX-304 treatment. Lean, fat and total body mass before (T=3w) and after (T=9w) ATX-304-treatment. **p<0.01,***p<0.001, ****p<0.0001 (One-way ANOVA with Tukey's multiple comparisons test). Individual data points, mean ± SEM are indicated (n=10 for all groups).

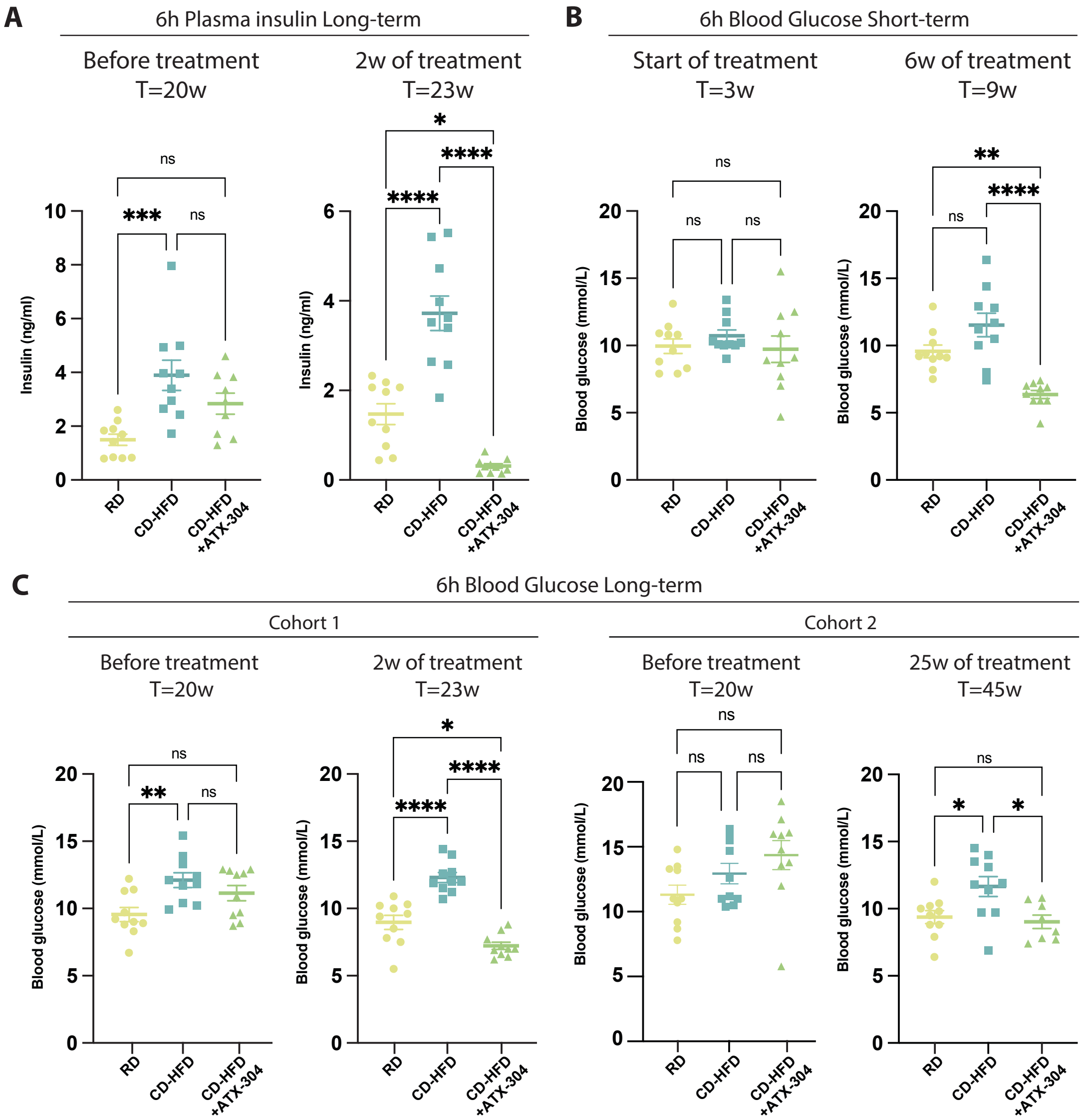


**Supplementary Figure 2. ATX-304 treatment significantly lowers insulin and blood glucose in C57Bl/6J male mice. A**) Fasted insulin for RD, CD-HFD and CD-HFD+ATX-304 before start (T=20 weeks) and after 2w (T=23 weeks) of O304 treatment. **B**) Fasted blood glucose for RD, CD-HFD and CD-HFD+ATX-304 at start (T=3 weeks) and after 6w (T=9 weeks) of ATX-304 treatment. **C**) Fasted blood glucose for RD, CD-HFD and CD-HFD+ATX-304 at start (T=20 weeks) and after 2w (T=23) or 25w (T=45w) of ATX-304 treatment. *p<0.05, ***p<0.001, ****p<0.0001 (One-way ANOVA with Tukey's multiple comparisons test). Individual data points, mean ± SEM are indicated in all graphs (n=10 for all groups).

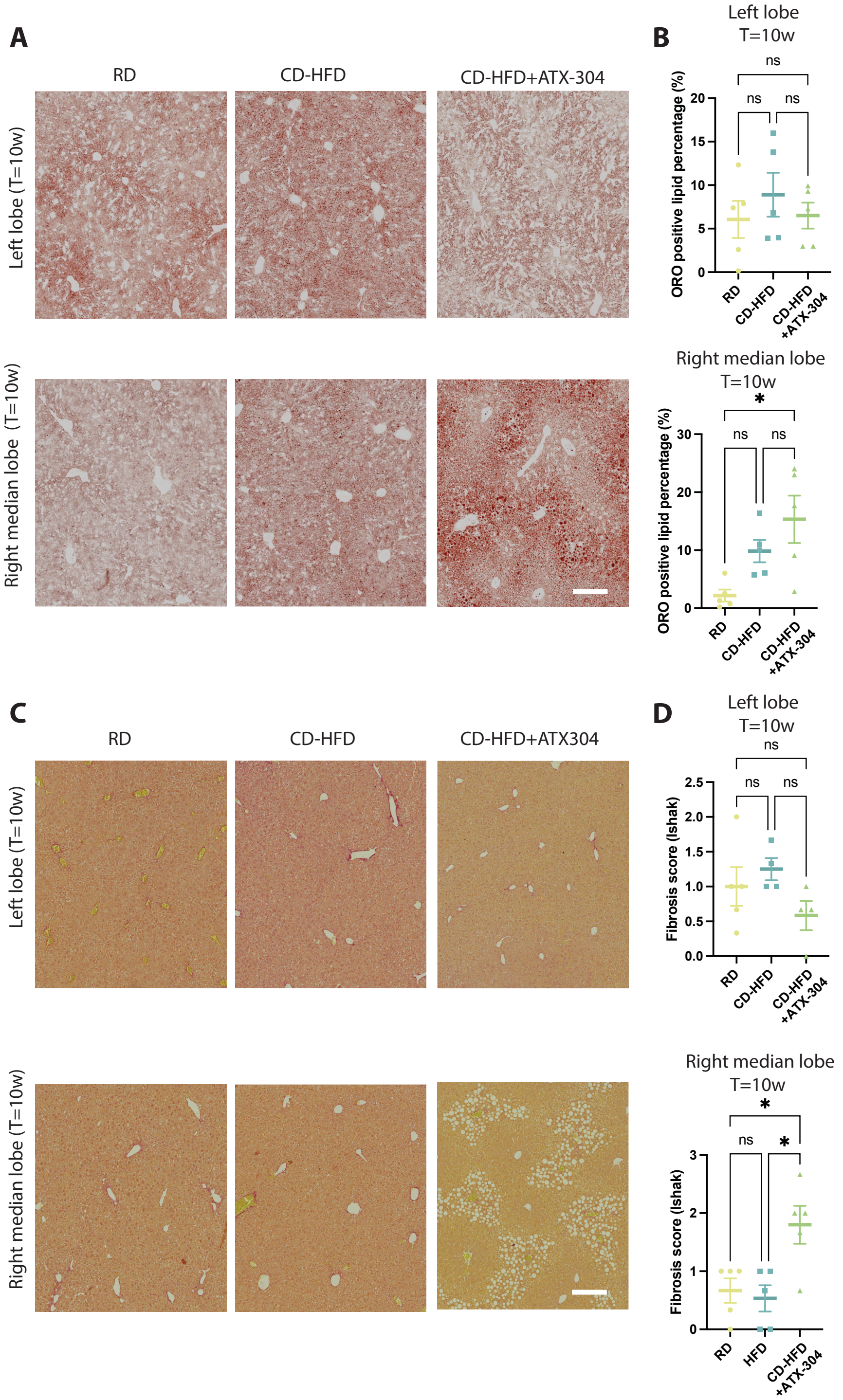


**Supplementary Figure 3. Lobular heterogeneities in distribution of lipids and fibrosis in ATX-304-treated livers. A**) Oil redO stained liver sections from the left lobe (upper row) and the right median lobe (bottom row) of RD, CD-HFD and ATX-304 treated mice at T=10w. Scalebar corresponds to 200um. Percentage of ORO positive area for the left (top) and right median lobe (bottom) is displayed in **B**) (n=5 for all groups). **C**) Representative images depicting picrosirius red (PSR) staining of sections from the left lobe (top row) and right median lobe (bottom row) from RD, CD-HFD and CD-HFD+ATX-304 livers at T=10w. Scalebar corresponds to 200um. **D**) Fibrosis score based on PSR staining for the left (top) and right median lobe (bottom). *p<0.05, ***p<0.001, ****p<0.0001 (One-way ANOVA with Tukey's multiple comparisons test). Individual data points, mean ± SEM are indicated in all graphs (n=5 for RD and CD-HFD, and n=4 for CD-HFD+ATX-304).

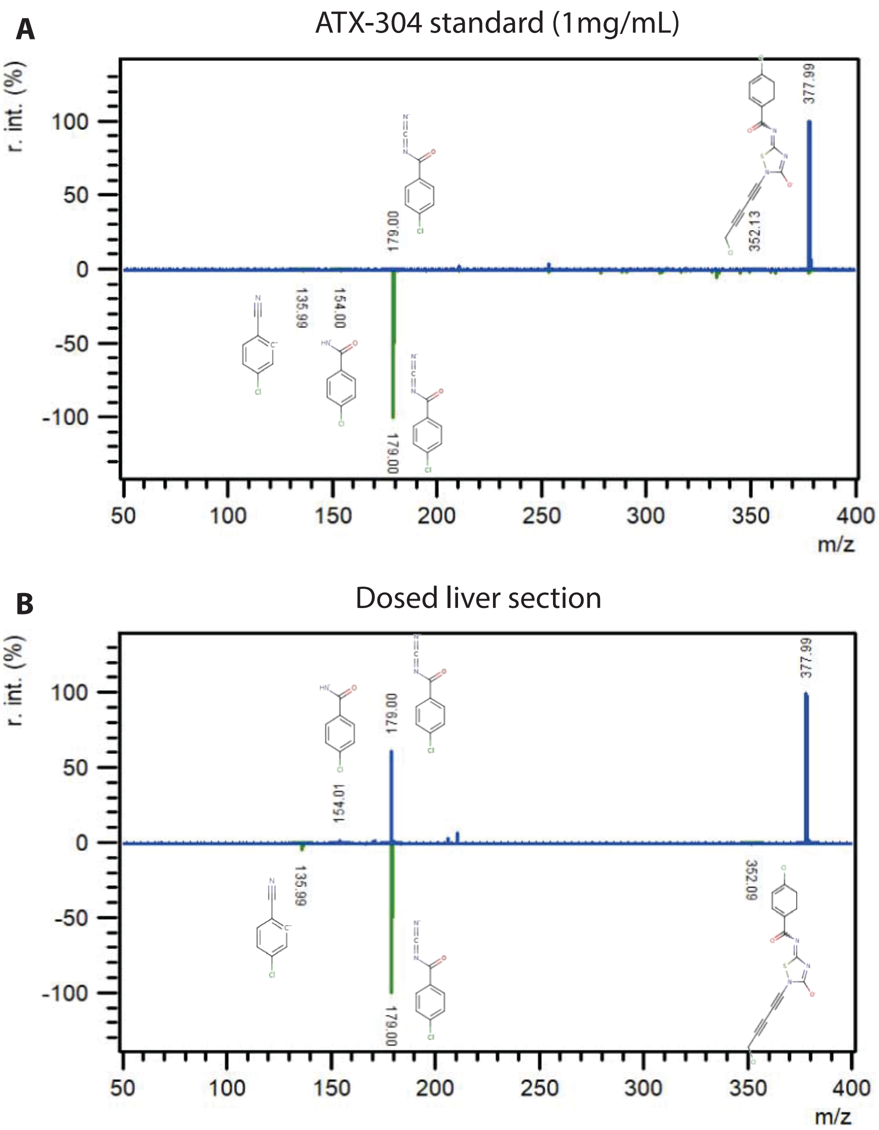


**Supplementary Figure 4. Identification of ATX-304 MS/MS spectrum in liver tissue sections.** MS/MS spectra from **A)** ATX-304 drug standard (1mg/mL) spotted on ITO slide and **B**) dosed liver section.
